# Habitat isolation diminishes potential of self-organised pattern formation to promote local diversity in metacommunities

**DOI:** 10.1101/2024.02.22.581536

**Authors:** Louica Philipp, Toni Klauschies, Christian Guill

## Abstract

Progressive destruction and isolation of natural habitat is a major threat to biodiversity worldwide. In this study we use a trophic metacommunity model with complex, spatially explicit structure to address how the interaction of local and regional processes affects the functional diversity of autotroph (producer) communities within and between individual habitat patches. One important driver of biodiversity in metacommunities is spatial heterogeneity of the environment, as it enables source-sink dynamics between patches. Besides a-priori differences in the environmental conditions, heterogeneous distributions of resources and species biomasses can also emerge through self-organised pattern formation caused by scale-dependent feedback between local trophic and regional dispersal dynamics. We show that this emergent heterogeneity can enhance the functional diversity of local autotroph communities by jointly strengthening source-sink dynamics and reducing stabilising selection pressure. Our results indicate that this effect is particularly strong in highly connected metacommunities, while metacommunity size (number of patches) alone plays a lesser role. We demonstrate that the positive effect on local diversity is driven by an eco-evo-spatial feedback loop that is fueled by the asynchronous biomass- and trait dynamics between the patches created by self-organised pattern formation. In highly connected metacommunities, oscillatory biomass patterns with particularly large amplitude strengthen this feedback loop. Our findings are highly relevant in the light of anthropogenic habitat changes that often destroy dispersal pathways, thereby increasing habitat isolation, lowering overall connectance of metacommunities and ultimately threatening the biodiversity in local habitats. Only a joint investigation of the contributing ecological, evolutionary, and spatial mechanisms in complex model systems can yield comprehensive understanding of these processes, allowing for the development of strategies to mitigate adverse anthropogenic influence.

## 1 Introduction

Agriculture, settlements and other human land uses are increasingly fragmenting natural landscapes into separate habitat patches. From an ecological perspective, these habitats are linked through the dispersal of species into large-scale metacommunities, which are often characterised by irregular spatial structures and complex dynamics, and they might still maintain high biodiversity (D. L. Urban and Keitt, 2001; Leibold et al., 2004; Holyoak et al., 2005; Holland and Hastings, 2008; Riva et al., 2023). The spatial structure of metacommunities, i.e., the number of habitat patches and dispersal pathways (’links’) between them (Fig. 1a) is indeed an important determinant for their diversity, as it controls the strength of dispersal-driven mass-effects. While stabilising selection and competitive exclusion can reduce the diversity within habitats (Norberg et al., 2001; Leibold et al., 2004), a moderate dispersal rate between adjacent habitats can support higher diversity through source-sink dynamics and spatial insurance effects (Brown and Kodric-Brown, 1977; Mouquet and Loreau, 2003; Cadotte, 2006; Staddon et al., 2010), especially when environmental conditions and thus local species communities differ between the habitats (Tews et al., 2004; Haegeman and Loreau, 2014). Conversely, if dispersal rates are too high, local and regional diversity might be lost through regional genetic or phenotypic homogenisation of local communities (Cadotte, 2006).

**Figure 1.**
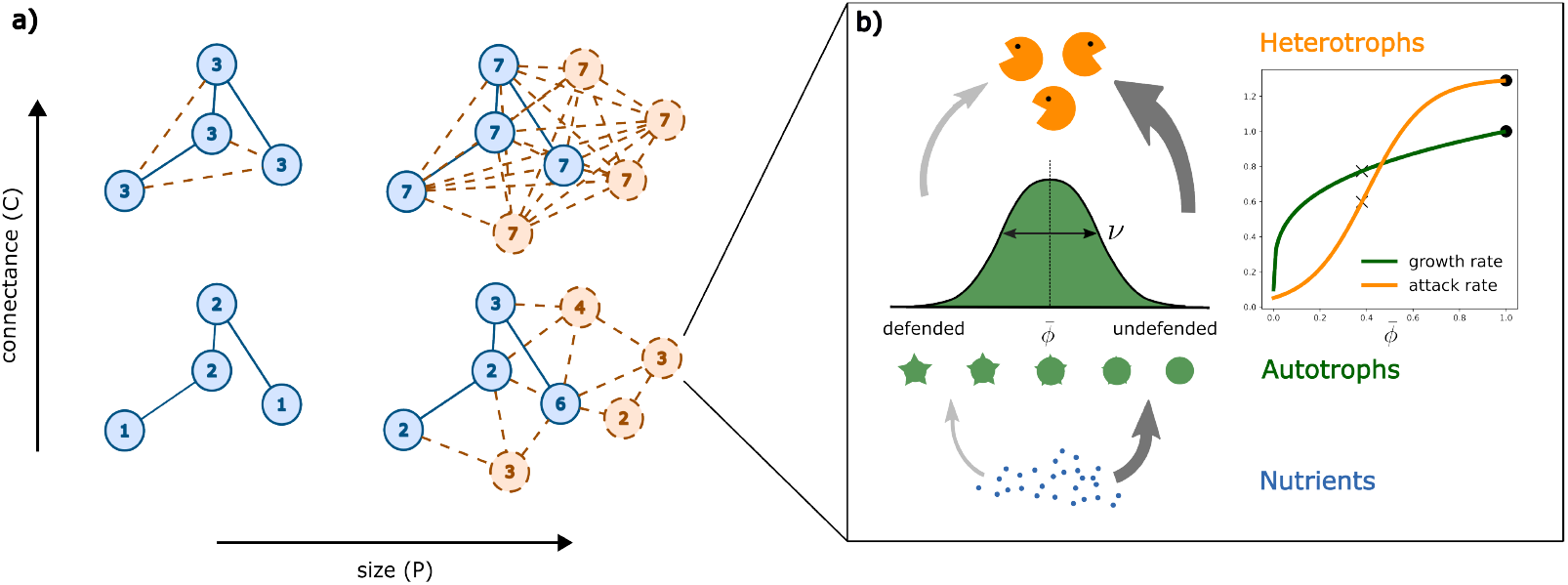
: a) Habitat change can alter the size and connectance of a metacommunity as dispersal links are added or deleted (vertical axis) or habitat patches, including their dispersal links, (dis)appear (horizontal axis). Numbers denote patch degrees, i.e. the number of dispersal links to other habitats. b) Aggregated food-web within a local habitat and trade-off between intrinsic growth rate of the autotrophs and attack rate of the heterotrophs. Both rates are dependent on the autotroph trait *ϕ* that has a logit-normal distribution in the local communities. The latter is described by mean trait 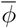 and trait variance *v*. Cross markers denote growth and attack rates at the defended, dot markers at the undefended local attractor.

The progressive intensification of human land use changes these spatial structures (Chase et al., 2020) and is a major threat to the biodiversity worldwide (Foley et al., 2005; Newbold et al., 2015; Balvanera et al., 2019). Through disruption of dispersal pathways or destruction of adjacent habitat patches, isolation of the remaining patches increases and mass-effects vanish (Staddon et al., 2010; Thompson et al., 2017; Ryser et al., 2019; Chase et al., 2020). Diversity thus decreases as single species or functional groups dominate in isolated communities (LeCraw et al., 2014). In degrading landscapes, the decrease of habitat connectance was found to be a stronger driver for loss of species diversity than reduced habitat size and quality itself (Horváth et al., 2019). Furthermore, processes on the local and regional scale are often interdependent and interact (Logue et al., 2011), which can lead to the emergence of complex dynamics on the metacommunity scale. Viewing (meta-)ecosystems as complex systems can help to understand potential effects of global change and is thus becoming increasingly relevant in scientific literature (Bauer et al., 2021; Riva et al., 2023). It is however not well established how complex dynamics and feedback mechanisms in metacommunities affect their diversity and how these effects are dependent on the spatial structure. A complex, dispersal-driven phenomenon observed in many different ecological systems is the formation of self-organised spatial patterns, where spots, stripes and other regular patterns in the local resource availability and biomass densities of species are formed on the regional scale (Malchow, 1993; Rietkerk and van de Koppel, 2008; Meron, 2015). This happens due to an interaction between the local dynamics and unequal dispersal speeds of resources (water, nutrients) and their consumers, which causes local accumulation and lateral depletion through scale-dependent-feedback (Murray, 2003; Rietkerk and van de Koppel, 2008; Kéfi et al., 2010). Most extensively studied are the self-organised vegetation patterns in arid ecosystems and savannas (Rietkerk and van de Koppel, 2008; Kéfi et al., 2010; Meron, 2015), but pattern formation is also known from a variety of other ecosystems, such as wetlands (van der Valk and Warner, 2009), intertidal mudflats (Liu et al., 2014) and aquatic communities (Cornacchia et al., 2018).

Recently, several studies have shown that self-organised pattern formation can contribute to the diversity of ecological systems. Since its first general description by Turing (1952) the phenomenon was traditionally analysed in continuous space, i.e., uniform landscapes (Murray, 2003; Meron, 2015). There, the formation of a spatial pattern can promote coexistence between (autotrophic) consumer species that differ in their dispersal strategies (Nathan et al., 2013; Eigentler and Sherratt, 2020) or through relaxing competition and even enabling facilitation between species in the emergent heterogeneous environmental conditions (Cornacchia et al., 2018; L. Eigentler, 2021; Bennett et al., 2023). These studies however neglect the fragmentation of many natural landscapes into discrete habitat patches that may be only partially connected by dispersal links and therefore often have irregular spatial structures (Holland and Hastings, 2008).

Self-organised pattern formation is also studied in spatially discrete, network-organised systems such as trophic metacommunities (e.g. Othmer and Scriven, 1971; Nakao and Mikhailov, 2010; Brechtel et al., 2018; Guill et al., 2021). In a metacommunity with six habitat patches arranged into a ring, Guill et al. (2021) showed that the local functional diversity of autotroph communities characterised by a continuous trait distribution (Fig. 1b) can be enhanced by pattern formation beyond the effect expected from source-sink dynamics alone. They hypothesised that this is possible as the self-organisation provides an emergent heterogeneity in the available nutrients and predator density in space and time, thereby leading to locally and temporally differing selection pressures and thus differentiated niches, to which the trait distributions of local communities keep adapting over time. Immigration of maladapted specimen from adjacent communities with different trait distributions also widens the local trait range and thus enhances the local trait diversity of the autotroph communities.

For network-organised systems such as metacommunities, it is recognised that the number of habitat patches and the specific arrangement of dispersal links determine if patterns form and what they look like (Box 1, Brechtel et al., 2018; Nakao and Mikhailov, 2010; Aufderheide, 2014; Van Der Kolk et al., 2023). Therefore, land use change that alters the spatial structure of a metacommunity might affect self-organised pattern formation, and in consequence threaten the diversity of local communities. In this study we thus expand the trophic metacommunity model of Guill et al. (2021) to assess the impact of two structural properties directly affected by anthropogenic changes, the number of habitat patches and dispersal links, i.e., the size and connectance, of a metacommunity, on the functional diversity within and between autotroph communities (Fig. 1). We expect that pattern formation will occur in metacommunities of various irregular structures and sizes (although for varying dispersal conditions, see Box 1) and consequently result in heterogeneous environmental conditions between habitat patches. We hypothesise that this creates source-sink dynamics between autotroph communities that will be stronger in metacommunitites with more dispersal links, i.e., in large or highly connected metacommunities (Fig. 1a). Maintenance of local diversity might thus be enhanced compared to small or sparsely connected systems. Simultaneously, the risk for regional homogenisation between the habitats could be increased as well due to higher average dispersal rates. However, larger metacommunities might also show spatial clustering of patches well connected to each other, but with fewer links to other clusters. As this could counteract homogenisation on the regional scale it remains open whether positive effects of enhanced source-sink dynamics or negative effects due to homogenisation dominate, or whether interactions occur. Lastly, changes to size and connectance of the metacommunity might alter the emerging patterns themselves. As pattern formation determines the redistribution of nutrients and species’ biomass in complex ways in space and time, any alteration of the pattern would again affect local diversity through source-sink dynamics, but may also affect local dynamics through changing the relative importance of top-down and bottom-up control in the food chain. As the modification of the patterns in metacommunities of different size and connectance might not follow a linear trend, local and regional interactions as well as diversity could change in non-intuitive ways.

### Box 1.

Self-organised pattern formation in network-organised metacommunities

Self-organised pattern formation is caused by a spatial (Turing) instability of a locally stable, homogeneous state. Whether such an instability occurs depends on the specific combination of diffusion coefficients *d*_*X*_ (*X* = *N, A, H*) and the metacommunity structure (given by its Laplacian matrix **L**, for example Fig. 2b) and can be analytically deduced from evaluation of the eigenvalue spectrum of a set of Turing matrices **T**_**i**_ given by

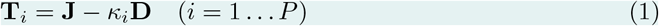

**Figure 2.**
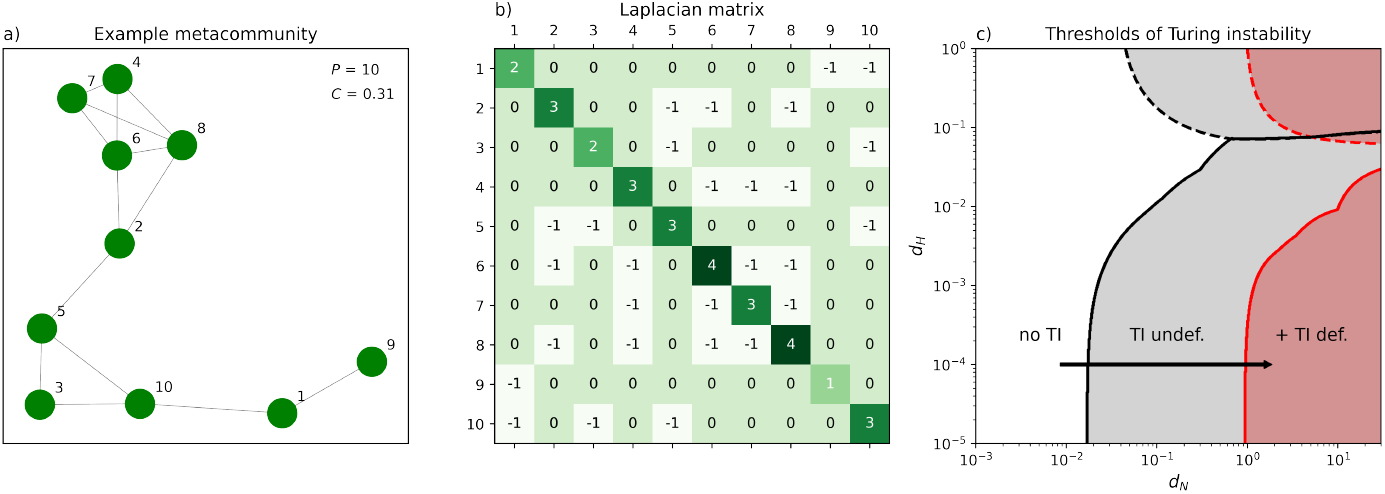
: a) Topology of an example metacommunity with *P* = 10 randomly placed habitat patches and 14 of 45 possible dispersal links connecting the patch pairs of shortest distance. Connectance *C* ≈ 0.31. b) Laplacian matrix **L** of the example metacommunity. Nondiagonal elements contain the adjacency information: *L*_*ik*_ = −1 if patch *i* and *k* are connected by a dispersal link and *L*_*ik*_ = 0 otherwise. Diagonal elements *L*_*ii*_ contain the patch degrees (sum of links to other patches). c) Regions in the nutrient and heterotroph diffusion coefficient space (autotroph diffusion coefficient *d*_*A*_ = 10^−5^) where self-organised pattern formation is predicted for the example metacommunity as identified from the leading eigenvalue *λ*_*max*_ of the Turing matrices **T**_*i*_ (Eq. 1). Areas with positive, real *λ*_*max*_ values (in dashed boundaries) indicate static patterns, positive, complex *λ*_*max*_ values (in solid boundaries) indicate oscillatory patterns. Grey areas: pattern formation of undefended autotroph communities, red areas: pattern formation of defended autotroph communities. The arrow denotes the gradient of *d*_*N*_ used in this analysis.

(Brechtel et al., 2018; Guill et al., 2021). Here **J** is the Jacobian matrix of the dispersing local food web components (*N, A, H*) evaluated at the homogeneous equilibrium, **D** is the Jacobian matrix of the emigration dynamics (which in the case of diffusive, i.e., random, dispersal is simply a diagonal matrix of the three diffusion coefficients *d*_*X*_) and *κ*_*i*_ are the eigenvalues of **L**, their number corresponding to the number of habitat patches (*P*). For a spatial instability it is sufficient that the leading eigenvalue *λ*_*max*_ across all *P* Turing matrices has a positive real part (Othmer and Scriven, 1971; Brechtel et al., 2018; Guill et al., 2021).

Emerging patterns are static (only spatial heterogeneity) if *λ*_*max*_ is a real number, and oscillatory (with spatio-temporal variation) if *λ*_*max*_ is a pair of complex conjugate numbers. Regions of static and oscillatory Turing instability in the *d*_*N*_ -*d*_*H*_ -plane evaluated from *λ*_*max*_ values are visualised for an example metacommunity in Fig. 2c.

The *κ* spectrum generally depends on the degrees (number of dispersal links) of the habitat patches (Hedetniemi et al., 2016). With changing size and connectance, but also between specific topologies of the metacommunities with the same spatial metrics, the patch degrees vary, thus altering the *κ* spectrum and the *d*_*X*_ thresholds for the Turing instability (Fig. 2c). For the combinations of *d*_*X*_ chosen in our study, higher patch degrees, which occur in larger and/or better connected metacommunities and result in larger *κ*, generally lowered the threshold of *d*_*N*_ necessary to induce pattern formation.

## 2 Methods

To simulate metacommunities with a wide range of spatial structures, we adapted the trophic metacommunity model from Guill et al. (2021). The food web within local habitats remained unchanged and its dynamics are described subsequently. Equations describing the metacommunity dynamics including dispersal between patches are given in section 2.2.

### 2.1 Local food web dynamics

Within each habitat patch we consider the dynamics of a small food web of nutrients with concentration *N*, a diverse autotroph community with total biomass density *A* and a heterotroph species with biomass density *H* (Fig. 1b). The food web is placed in a chemostat environment implemented with a fixed supply concentration of nutrients (*S*, identical on all patches) and a dilution rate (*D*) that affects all components equally. Nutrient uptake by the autotrophs follows Michaelis-Menten-kinetics, described by a half-saturation constant *N*_*H*_ and a maximum growth rate *r*(*ϕ*) that depends on a functional trait *ϕ*. Autotroph biomass is consumed by heterotrophs with grazing rate *g*(*ϕ*), which is given by a type II functional response with trait-dependent attack rate *a*(*ϕ*) and handling time *h*. Autotroph biomass acquisition and loss can be summarised into its net per-capita growth rate (fitness function) *G*_*A*_(*ϕ*). Heterotroph biomass growth from grazing is scaled with the conversion efficiency *e*. Taken together, the temporal change of nutrient concentration *N* and biomass densities *A* and *H* are given by the following differential equations:

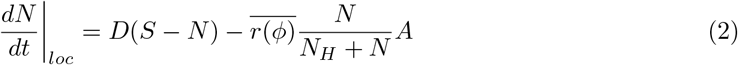

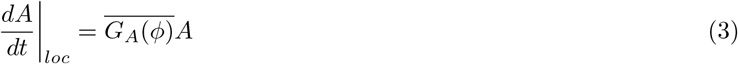

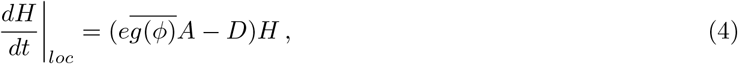

With

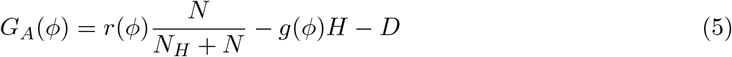

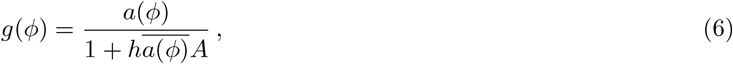

and trait-dependent growth rate and attack rate functions defined as

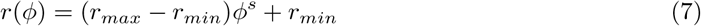

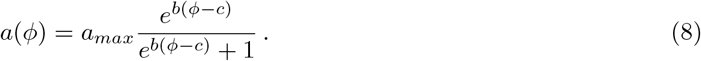

The growth rate *r*(*ϕ*) is a power law function within assumed lower and upper limits *r*_*min*_ and *r*_*max*_ (Fig. 1b). The curvature of the function is determined by the parameter *s*. The attack rate function describes a sigmoidal curve with maximum attack rate *a*_*max*_ (Fig. 1b). This saturating curve is shaped by sensitivity parameter *b* and inflection point *c*.

We assume the trait *ϕ* has variation in the local autotroph communities and occurs with a logit-normal distribution in the limits [0,1], where high values indicate high maximal growth rates of the autotrophs and high attack rates of the heterotrophs (Fig. 1b). Low trait values in turn are associated with lower attack rates (e.g. through a defence mechanism) but also lower intrinsic growth rates. This growth-defence trade-off determines the shape of the fitness landscape and thus the selection pressure at a particular moment in time (Tirok et al., 2011). It ensures that defended types (low *ϕ*) will thrive when heterotrophs are abundant, but are at a disadvantage compared to undefended types (high *ϕ*) when heterotroph biomass is low.

All functions that depend on the trait distribution are modelled as average functions 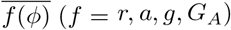. Following an aggregate modelling approach (Wirtz and Eckhardt, 1996, Norberg et al., 2001, Klauschies et al., 2018), these can be sufficiently described by second-order approximations using the trait distribution mean, 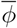, and variance, *v*, as

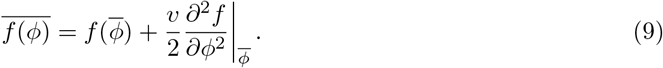

The temporal changes of mean and variance of the trait distribution in a local autotroph community are given as

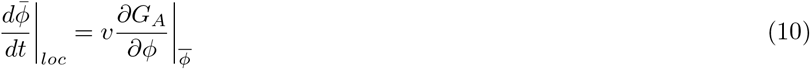

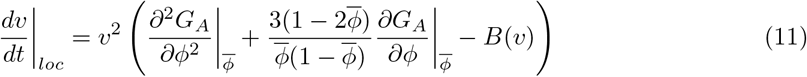

with a boundary function

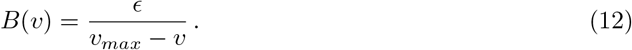

Changes in 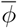 and *v* are driven by the local fitness gradient 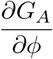 at 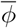 and the local shape of the fitness landscape 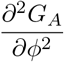 around 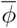 (Klauschies et al., 2018). Specifically, 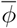 is changing in the direction that enhances the per capita net growth rate of the autotrophs. The speed of change is scaled by *v*, which reflects the functional diversity of the autotrophs: the presence of many different functional types (*v* large) enhances the capability of a community to adapt, while a lack of functional diversity (*v* small) slows down the adaptation process. 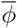 will stop changing when it reaches a local fitness maximum or when *v* is reduced to zero and thus the potential for adaptation is lost. Internal changes in *v* are predominantly determined by the local curvature of the fitness landscape, which accounts for stabilising or disruptive selection (reducing or enhancing *v*, respectively). The boundary function *B*(*v*) keeps *v* below a maximum value of *v*_*max*_. The scaling parameter *ϵ* is chosen small enough to not affect the model dynamics when *v* is further away from *v*_*max*_.

Modelled in isolation, the local food web dynamics are bistable (Fig. 1b). The autotroph community either converges towards a low trait value 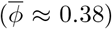, implying low growth rates but also a lower heterotroph attack rate (from here on called ‘defended’ attractor and community). Alternatively, the local autotroph community can settle onto a high trait value 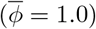 implying a high autotroph growth rate but also a high heterotroph attack rate (the ‘undefended’ attractor and community). On completely isolated patches and 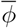 close to the respective attractor, *v* always slowly decays to zero due to stabilising selection, thereby preventing any further temporal change of the mean trait.

### 2.2 Metacommunity model

A copy of the aggregated food web is placed on each of *P* habitat patches in a two-dimensional landscape. The horizontal and vertical coordinates of each patch are randomly drawn from a uniform distribution within the limits [0, 1]. Undirected dispersal links connect the local habitats to metacommunity graphs (Fig. 1). The connectance *C* of a metacommunity is given as

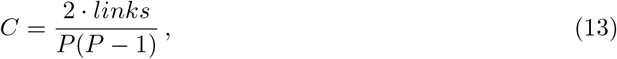

the ratio of the number of existing dispersal links divided by the maximum number of possible links in a metacommunity of size *P*, excluding self-links (Bornholdt and Schuster, 2001). To generate metacommunity graphs with a desired connectance, the required dispersal links are set between the pairs of patches with the shortest Euclidian distance. In the generation process we discarded metacommunities consisting of disconnected subgraphs. Adjacency information of a metacommunity graph is stored in the Laplacian matrix **L** of size *P × P*, where nondiagonal elements *L*_*ik*_ = −1 if patches *i* and *k* are connected and *L*_*ik*_ = 0 otherwise (for example see Fig. 2b). Diagonal elements *L*_*ii*_ are assigned the respective patch degree, i.e. the sum of dispersal links to other patches.

Metacommunity dynamics are determined by both the local interactions 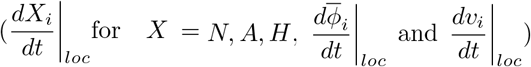 and the spatial dynamics, i.e., dispersal, and are for any patch *i* described by the following differential equations:

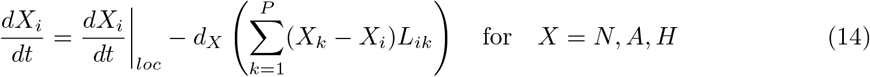

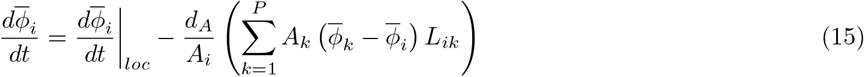

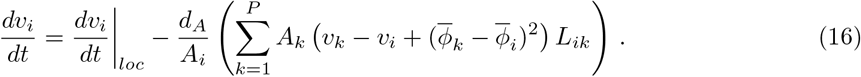

The dispersal of nutrients, autotrophs and heterotrophs between adjacent patches (*L*_*ik*_ = −1) is driven by differences in concentration or biomass density, respectively (diffusive movement) and is additionally scaled by a diffusion coefficient *d*_*X*_ (*X* = *N, A, H*). With the exchange of autotroph biomass (of potentially different trait distributions), the local mean traits 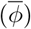 and variances (*v*) may also change (Norberg et al., 2001; Klauschies et al., 2018). The change of the resident mean trait value is affected by the difference between mean trait values of resident and incoming species as well as by the ratio of incoming to resident biomass. The change of the trait variance is affected in a similar way by the difference between trait variances of resident and incoming species, and it increases if the mean trait of the incoming species differs from that of the resident.

### 2.3 Functional diversity

We differentiate between the mean local (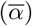) and between-patch (*β*) functional diversity of an autotroph metacommunity. The local functional diversity of the autotroph community on a given patch is given by the variance *v*_*i*_ of its local trait distribution. High variance implies a wide trait distribution and is thus associated with a large range of trait values expressed with reasonably high biomass density. Between-patch functional diversity is measured as the difference between the mean traits 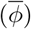 of the local communities, with larger differences between patch means associated with higher *β*-diversity. In line with Guill et al. (2021), the biomass-weighted average local 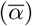 and between-patch (*β*) diversity of a metacommunity are given by

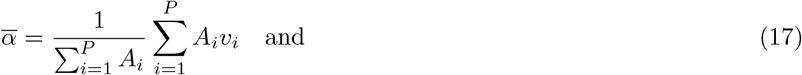

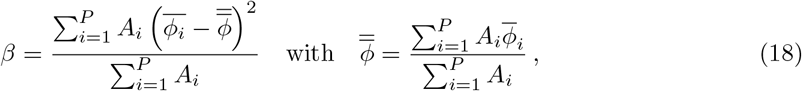

where 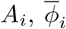 and *v*_*i*_ are temporal averages and 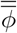 is the grand mean trait across all patches of a metacommunity. Regional diversity (*γ*) can be calculated as the sum of 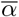 and *β* diversity (see supplementary material of Guill et al. (2021) for derivation) and is therefore not separately evaluated here.

### 2.4 Simulation protocol

Functional diversity was assessed from numerical simulations of the dynamics in randomly generated metacommunities of size *P* and connectance *C*. The nutrient diffusion coefficient *d*_*N*_ was systematically varied as a key metric determining whether pattern formation occurs for the undefended or defended attractor (see Fig. 2c). To limit the complexity of the analysis the remaining diffusion coefficients *d*_*A*_ and *d*_*H*_ were kept at a constant, low level chosen so that consistently oscillatory patterns emerged (figure 2c). The remaining model parameters were set to constant values oriented at planktonic systems (Appendix table A1).

To assess the effect of the metacommunity structure on functional diversity, three sets of simulations were run. In the first, metacommunities with a fixed size of *P* = 10 patches were modelled at 17 levels of connectance. Connectance was systematically varied from 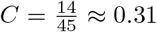 to C=1 by adding two links in each step (one in the last). The minimum connectance was chosen as especially in small metacommunities, connectance ⪅0.3 resulted in a high fraction of the randomly created graphs to be disconnected and therefore discarded. Nutrient mobility *d*_*N*_ was varied as a second gradient from 0.008 to 1 in 30 geometrically evenly spaced steps. At each connectance level, 100 metacommunities with randomly chosen patch positions were generated. For simulating metacommunity dynamics, initial values of all variables were randomly drawn with uniform probability from the intervals *N*_*i,init*_ = [0.1, *S*], *A*_*i,init*_ = [0.1, *S*], *H*_*i,init*_ = [0.1, *eS*], 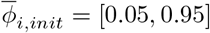, *v*_*i,init*_ = [0.005, 0.01].

For a second and third set of simulations, metacommunities of sizes *P* = 10 to *P* = 70 with an increment of 10 patches and fixed, respectively low (*C* = 0.3) or high (*C* = 1.0) connectance were generated. Due to shifted thresholds for the Turing instability in larger metacommunities, *d*_*N*_ was varied in an extended range from 0.001 to 1.1 in 30 geometric steps. For both sets, dynamics of 40 randomly generated metacommunities of each size were simulated with random initial conditions as specified above. As the simulation times for the dynamics of especially the large metacommunities were comparably high and the results showed only limited variability, we refrained from including more replicates in this part of the analysis.

All simulations were run for 10^6^ time steps to eliminate transient dynamics and then evaluated over another 1.5 · 10^4^ time steps. The model output consisted of long-term temporal averages and variances of the variables of each patch.

Simulations were written in the programming language C. Differential equations were numerically solved using the BDF method of the SUNDIALS CVODE solver version 6.1.1 (Hindmarsh et al., 2005) with relative and absolute tolerances of 10^−15^. Vector and matrix operations of the GNU scientific library (Galassi et al., 2002) were implemented to increase runtime efficiency. C simulations were coordinated and additional tasks as well as data analysis and visualisation were conducted in Python ver. 3.10.4 using packages NumPy ver. 1.22.3 (Harris et al., 2020), SciPy ver.

1.8.0 (Virtanen et al., 2020), Joblib ver. 1.1.0 (Joblib Development Team, 2021) and Matplotlib ver. 3.5.1 (Hunter, 2007).

## 3 Results

### General effects of source-sink dynamics and pattern formation on diversity

Independent of pattern formation, a low level of mean local functional diversity (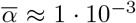 in smaller metacommunities and up to ≈ 7 · 10^−3^ in larger metacommunities, Figure 3 a, c, e) and a high level of between-patch diversity (*β* ≈ 7 · 10^−2^, Figure 3, b, d, f) is maintained whenever both undefended and defended attractors are present in a metacommunity. The exchange of autotroph biomass with respectively high and low mean traits 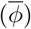 preserves local trait variances (*v*) and thus 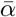-diversity at a nonzero level due to source-sink dynamics.

**Figure 3.**
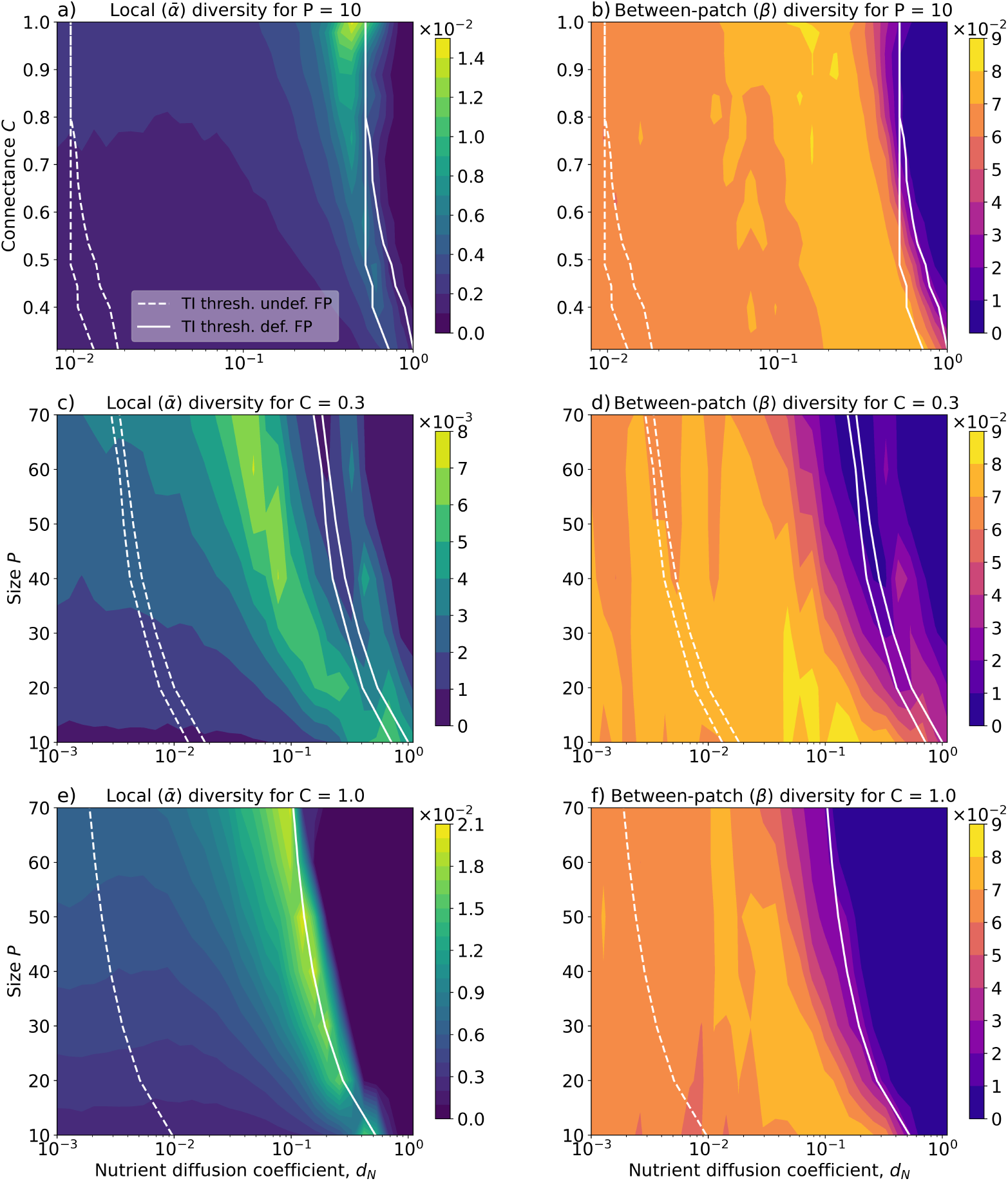
: Mean local (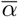) and between-patch (*β*) functional diversity in randomly generated metacommunities with size (number of patches) *P* and connectance (fraction of dispersal links) *C*. In panels a) and b), connectance is varied in metacommunities with a fixed size of *P* = 10. In panels c) - f), a gradient of metacommunity size is evaluated for low (0.3) and full (1.0) connectance, respectively. Note the different scales for 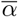-diversity between a), c) and e). In all panels, *d*_*A*_ = 10^−5^ and *d*_*H*_ = 10^−4^. White lines denote spatial instability thresholds of the undefended attractor (dashed) and the defended attractor (solid) predicted from Turing matrix eigenvalues (Box 1). For *d*_*N*_ above the thresholds, oscillatory patterns emerge. For both attractors only the lowest and highest threshold of the metacommunities used in simulations are shown. The range of instability thresholds reflects stochastic differences in the spatial structure of the metacommunities, which are largest for low *C* and disappear when *C* approaches 1.

With pattern formation, 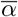-diversity can strongly increase above the source-sink level. Oscillatory pattern formation is induced for values of the nutrient diffusion coefficient *d*_*N*_ larger than a threshold value, which differs for the two attractors (Box 1). The amplitude of the oscillations subsequently increases over the *d*_*N*_ gradient (Supporting Information, Fig. A1). Enhancement of 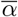 -diversity is especially pronounced for high *d*_*N*_ above the spatial instability threshold of the undefended attractor (dashed white lines in Fig. 3) and in proximity to the instability threshold of the defended attractor (solid white lines). Under these conditions especially undefended autotroph communities exhibit a strong oscillatory pattern in their biomasses, but defended autotroph communities already co-oscillate (Fig. 4, top left panels). We found a slight increase of *β*-diversity under pattern formation of the undefended communities for lower to intermediate *d*_*N*_ (Figure 3b, d, f). For high *d*_*N*_ in proximity to the instability threshold of the defended attractor, *β*-diversity however substantially decreases. Lowered levels of *β*-diversity coincide with maximum 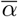-diversity. Eventually, a complete loss of both 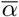- and *β*-diversity occurs when *d*_*N*_ is so high that the defended state becomes spatially unstable as well (beyond solid white lines in Fig. 3). We found that this results from homogenisation in the metacommunities, where all patches converge on a common (defended or undefended) state rather than maintaining two alternative stable states (details discussed in Supporting Information, Appendix A.3).

**Figure 4.**
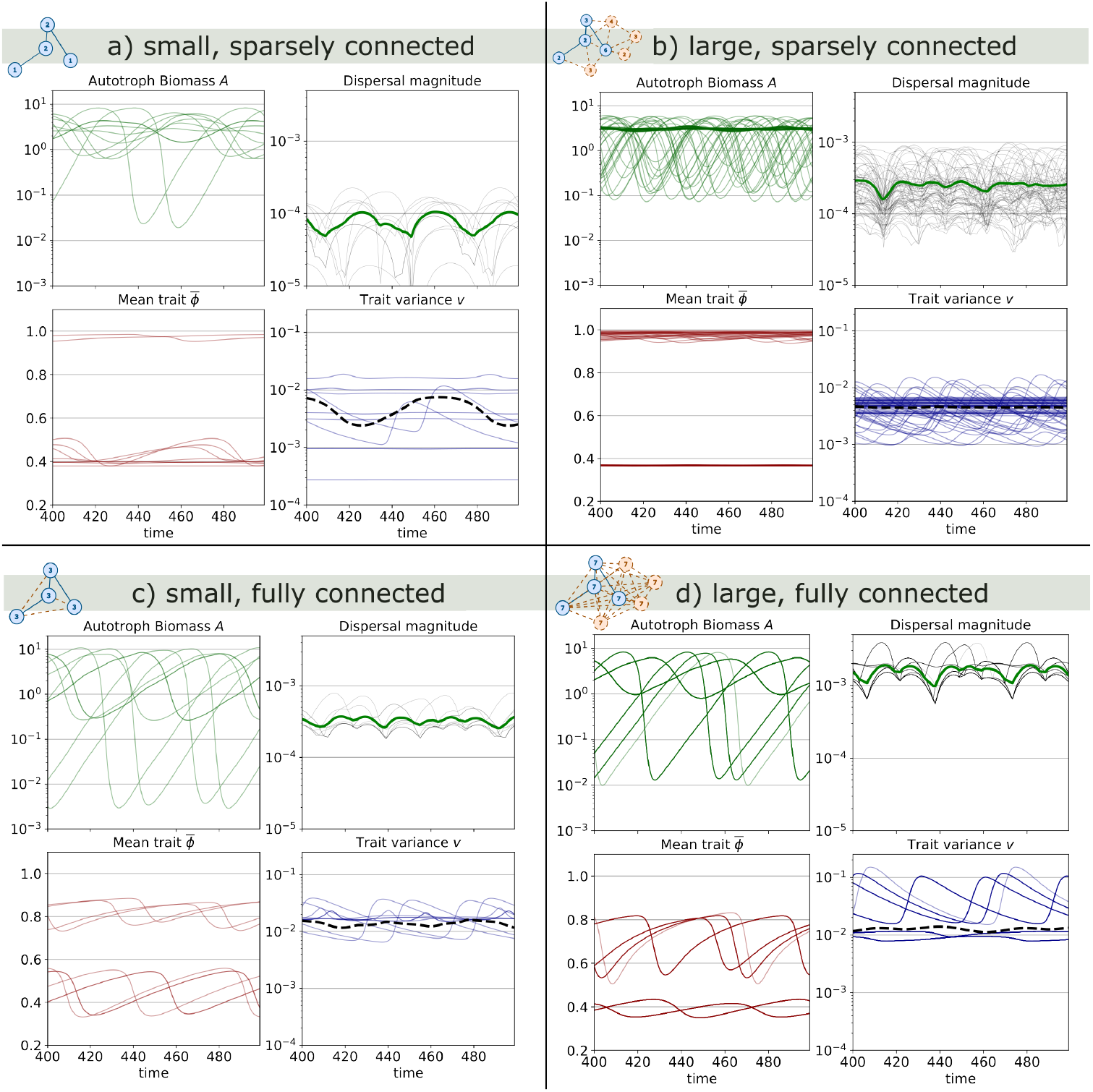
: Dynamics of autotroph biomass *A*, magnitude of autotroph dispersal per patch, trait distribution mean 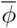 and variance *v* for a a) small, sparsely connected (*P* = 10, *C* = 0.3), b) large, sparsely connected (*P* = 70, *C* = 0.3), c) small, fully connected (*P* = 10, *C* = 1.0) and d) large, fully connected (*P* = 70, *C* = 1.0) metacommunity at their respective 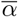-diversity peak. (a): *d*_*N*_ = 0.75, b): *d*_*N*_ = 0.05, c): *d*_*N*_ = 0.5, d): *d*_*N*_ = 0.09). The dispersal magnitude was calculated for all patches *i* as Σ*d*_*A*_(|*A*_*i,t*_ − *A*_*j,t*_|) (black lines) and averaged across all *P* patches (green line). Average *v* across patches is shown as dashed black lines. Dynamics were captured after 100000 transient time steps. Transparency of lines to visualise synchrony of dynamics.

The peak of 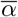-diversity roughly follows the instability threshold of the defended attractor, which occurs at lower *d*_*N*_ the higher connectance (*C*) or size (*P*) are. At low connectance (*C* = 0.3), the 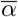-diversity peak is broad (Fig. 3 a, c), reflecting the stochastic differences in the spatial structures between metacommunities (Box 1). These however fade for high connectance (*C* ⪆ 0.7) and therefore instability thresholds as well as 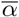-diversity peaks coincide between metacommunities (Fig. 3a, e). However, a similar effect does not occur for larger metacommunity size.

### Effects of metacommunity connectance on diversity

The peak value of 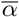-diversity developing under pattern formation increases with connectance of the metacommunities (Fig. 3a). Peak 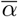-diversity increases 4-fold from sparsely connected (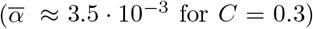 for C=0.3) to fully connected (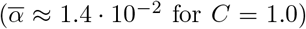 for C=1.0) small metacommunities (*P* = 10). At full connectance, 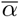-diversity increases 12-fold compared to the source-sink level, while for sparsely connected metacommunities, it only increases by a factor of 3. Complementing these results, peak 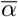-diversity is generally higher in fully connected than in sparsely connected metacommunities across different metacommunity sizes (compare Fig. 3c to e). We found that increasing connectance also marginally enhances *β*-diversity for intermediate levels of *d*_*N*_ (Fig. 3b). To illustrate the effect of connectance on the emerging pattern itself, we compare example dynamics of a sparsely connected (*C* = 0.3) against a fully connected (*C* = 1.0) 10-patch meta-community at their respective 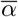-diversity peak (Fig. 4a, c). In both metacommunities the spatial instability of the undefended attractor causes asynchronous oscillations in the biomass of undefended, but also defended local autotroph communities over several orders of magnitude. The oscillation amplitudes however are visibly larger in the fully connected metacommunity. This stronger pattern yields larger momentary differences between the autotroph biomasses of adjacent patches, which result in higher dispersal flows per link. Additionally, the fully connected metacommunity has more dispersal links per patch (higher degrees, Fig. 1a) than the sparsely connected metacommunity. The metacommunity with higher connectance is thus characterised by a larger dispersal magnitude per patch (Fig. 4a, c). This higher dispersal flow correlates with the increasing peak 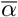 -diversity over the connectance gradient. A similar increase in pattern strength (amplitude of biomass oscillations), dispersal flow and peak 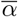-diversity from low to full connectance can be observed in the dynamics of large (*P* = 70) metacommunities (compare Fig. 4b to d). In the fully connected metacommunities, both undefended and defended communities show high *α*-diversity, i.e., variance of their local trait distributions *v* (Fig. 4c, d). High *v* occurs on all patches of the fully connected metacommunities and maintains flexibility of the local trait means 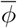 (see Eq. 10), which visibly oscillate as well. In the sparsely connected metacommunity, this is only possible on a few patches with favourable dispersal links to patches where the autotroph community exhibits a different attractor.

High autotroph dispersal flows lead to convergence of the 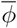 between the undefended and defended communities at the 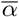-diversity peak in the fully connected metacommunities. The convergence of 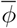 implicates a reduction of *β*-diversity prior to homogenisation within metacommunities (Supporting Information, Appendix A.3). In the sparsely connected metacommunities, oscillations and convergence of the local 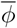 is only marginally visible. Eventually, *β*-diversity there is lost due to homogenisation within metacommunities as well.

### Effects of metacommunity size on diversity

With a larger metacommunity size, 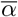 -diversity increases as well (Fig. 3c, e), albeit not as strongly as over the connectance gradient. In sparsely connected metacommunities, the overall highest 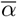 - diversity is 7 · 10^−3^, while in fully connected metacommunities it is 2.0 · 10^−2^. In both cases, these peak values are already reached in medium-sized metacommunities (*P* ⪆ 40). Further increase in the number of patches only widens the peak over the *d*_*N*_ -gradient, but does not increase its height. We also note that the level of 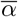-diversity that is solely maintained by source-sink dynamics (i.e., irrespective of self-organised pattern formation) increases with metacommunity size, too. This means that the relative increase of 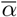-diversity caused by strong pattern formation is in large metacommunities not as pronounced as in small ones.

An increase in size slightly decreases the level of maintained *β*-diversity over much of the *d*_*N*_ range in both sparsely and fully connected metacommunities (Fig. 3d, f). Interestingly, in medium to large (*P* ⪆ 30), sparsely connected metacommunities (Fig. 3d) a secondary diversity peak emerges beyond the instability threshold of the defended attractor (where we usually observe homogenisation within metacommunities). At these high *d*_*N*_ values many metacommunities again exhibit local bistability and thereby maintain *β*-diversity. There, source-sink dynamics, enhanced by strong pattern formation, also lead to a secondary peak of 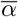 diversity (Fig. 3c). See also Supporting Information, Appendix A.3 for a more detailed discussion.

Examples of autotroph biomass dynamics of large metacommunities (Fig. 4b, d) show that the oscillation amplitudes at their respective 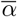-diversity peak in both cases are not higher than those of small metacommunities with equal connectance (Fig. 4a, c). While this implies that the dispersal flow per link does not change much between the small and large metacommunities, the greater number of links per patch in large metacommunities means that the absolute dispersal flow per patch is substantially higher in large than in small metacommunities (Fig. 4a to b, c to d). Note that in the large, fully connected metacommunity shown here, groups of patches with synchronised dynamics occur and dispersal is thereby limited to a subset of patches with asynchronous dynamics. Even though dispersal flow per patch is increased with size, this alone does not enable substantial fluctuations of the mean traits in the large, sparsely connected metacommunity (Fig. 4b).

### Role of local selection for the maintenance of local diversity

In general, the temporal dynamics of the trait variance *v* and thus the functional diversity of the autotrophic communities are influenced by both dispersal among different habitats and the respective local selection regime. While local selection mostly acts stabilising, thereby reducing *v*, dispersal always increases *v* through source-sink dynamics in our metacommunity model (Fig. 5). Under oscillatory pattern formation of the undefended local communities, local biomass density of the autotrophs strongly varies over time, and thus the strength of the dispersal effect on *v* changes over time as well, benefiting a local autotroph community more when it acts as a sink patch holding comparably low biomass. The magnitude (and sometimes even the direction) of local selection also strongly varies over time. In the undefended communities, stabilising selection is strongest when *v* peaks, as the trait distribution mean, 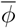, then is able to stay close to its optimum even under rapidly changing nutrient availability and predation pressure from oscillatory pattern formation.

**Figure 5.**
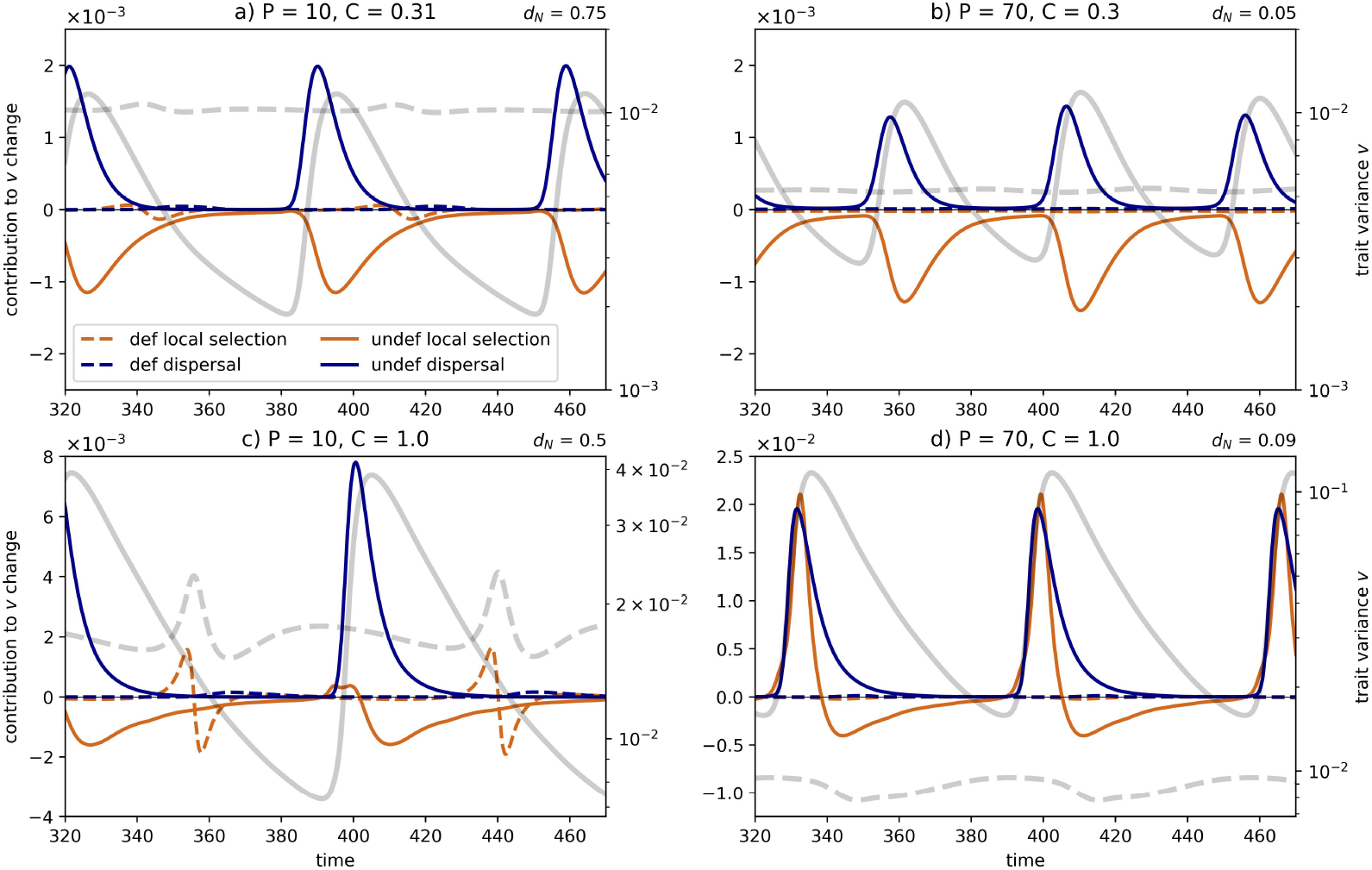
: Effect of local selection (orange, first term of the right-hand side of Eq. (16)) and dispersal (blue, second term of the right-hand side of Eq. (16)) on the dynamics of the trait variance (*v*) for the same example metacommunities as in Fig. 4. Each panel shows dynamics of one representative defended (dashed lines) and undefended (solid lines) community of the respective metacommunity. The dynamics of *v* are shown in grey as reference (scale on secondary vertical axis).

In consequence, *v* is again reduced (self-regulation of *v*). Low *v* however prevents adaptation of the mean trait to changing conditions, which implies less stabilising selection when the trait mean is further away from its optimum. In consequence, *v* can increase again through the positive influence of source-sink dynamics. The effects of dispersal and local selection on *v* are much less pronounced in defended communities, which also do not exhibit pattern formation in the shown examples (Fig. 5). Local diversity on these patches can however often be maintained at comparable levels.

In fully connected metacommunities, both small and large, the positive effect of dispersal on *v* is more pronounced, thus supporting maintenance of higher *v* levels. Additionally, the effect of local selection is temporarily positive, implying disruptive selection, which lets *v* increase even further. Disruptive local selection is especially pronounced in the large, fully connected metacommunity, where local selection at times increases variance as much as dispersal. Temporary disruptive selection also occurs in some of the defended local communities, which then also display high trait variance (best visible in Fig. 5 c). Hence, in addition to dispersal, the local selection regime is also affected by pattern formation and can substantially contribute to the maintenance of heightened local trait variance (*v*), i.e., local functional diversity, in the autotroph communities.

## 4 Discussion

In this study, we show that self-organised pattern formation can enhance the mean local 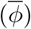 functional diversity in metacommunities with realistic spatial structure. Furthermore, we found that local diversity under pattern formation is especially pronounced in metacommunities with high connectance (number of dispersal links), while a large size (number of habitat patches) alone did only yield slightly higher local diversity. This differentiation can be explained by two distinct mechanisms: Firstly, habitat patches in both large and well-connected metacommunities have more dispersal links (i.e., higher patch degrees) and thus experience stronger source-sink dynamics, which results in the maintenance of a higher base-level local diversity. Secondly, highly connected metacommunities additionally exhibit patterns with larger oscillation amplitudes. This enhances the dispersal flow via existing dispersal links while simultaneously allowing for less stabilising or even disruptive local selection, which additionally contributes to the maintenance of higher local diversity. Especially in highly connected metacommunities, high local diversity of the autotroph communities enables continuous adaptation of their trait distributions to the fluctuations in nutrient availability and predation pressure, which again has a positive effect on local diversity through source-sink dynamics. This highlights an emergent eco-evolutionary feedback in our spatially explicit metacommunities with potentially far-reaching consequences for the adaptability and persistence of natural communities facing anthropogenic land use change.

### 4.1 Local diversity mechanism (1): Number of dispersal links

Typical patch degrees (i.e., number of dispersal links per patch) are higher in large and/or highly connected metacommunities than in small and/or sparsely connected ones (Fig. 1a). As hypothesised, large or well-connected metacommunities thus experience a higher net dispersal flow per patch (Fig. 4). Through coupling many autotroph communities with different trait distributions, they also exhibit stronger source-sink dynamics, which has a positive effect on the local trait variance, i.e., local functional diversity. This is in line with classic metacommunity theory, where a higher exchange of biomass between separated communities with different trait compositions is generally beneficial for local diversity (Amarasekare, 2003; Leibold et al., 2004; Haegeman and Loreau, 2014). This mechanism acts both with and without pattern formation as long as some between-patch (*β*) diversity is present in the metacommunity, which in our case is provided by the alternative attractors of the local communities. Therefore, local diversity in large or well-connected metacommunities increases from low to high nutrient diffusion coefficients (*d*_*N*_), including *d*_*N*_ values for which pattern formation is not induced. While the positive effect of metacommunity size on the local diversity maintained by source-sink dynamics is clearly visible (especially for low values of *d*_*N*_, Fig. 3c, e), the base-level local diversity is only slightly higher in well-connected metacommunities compared to sparsely connected ones (Fig. 3a). This is reasonable as for the size range considered in this study the patch degrees differ much more between small and large metacommunities than between sparsely and fully connected ones (Eq. 13 and Supporting Information, Figs. A2 and A3).

### 4.2 Local diversity mechanism (2): Strength of pattern formation

In metacommunities with high connectance local diversity is maintained at substantially higher levels through a second mechanism that directly depends on the occurrence of pattern formation. The amplitude of autotroph biomass oscillations, i.e., the ‘strength’ of the emergent oscillatory patterns, is more strongly affected by the connectance than by the size of metacommunities (Fig. 4 and Supporting Information, Fig. A1). Preliminary analyses suggest that this difference could be due to dissimilar changes of the underlying degree distributions in the metacommunities when size and connectance are modified (detailed in Supporting Information, Appendix A.2). Stronger oscillatory patterns imply higher momentary differences between the autotroph biomass densities of connected patches, which increases the net dispersal flow through individual dispersal links. Together with the higher total number of links in highly connected metacommunities as discussed above, this strongly enhances overall dispersal flow and thus source-sink dynamics between different autotroph communities, thereby allowing for the maintenance of high levels of local diversity.

The pattern strength additionally affects local diversity through modification of the local selection regime. The large amplitude oscillations of the nutrients, autotroph and heterotroph biomasses that result from the oscillatory Turing instability promote strong temporal fluctuations in the local optimal defence levels, which might strongly differ from the momentary realised mean trait 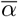 of the local autotroph communities. Consequently, this leads to less stabilising or even disruptive local selection on the local trait composition, which allows for the maintenance of higher trait variances (*v*) under pattern formation (already hypothesised by Guill et al., 2021), that directly translates to higher local diversity in the autotroph communities. High metacommunity connectance thus plays a twofold role: it allows for disruptive local selection while simultaneously leading to higher dispersal flow of autotroph biomass between habitats without inducing homogenisation, as observed when the nutrient diffusion coefficient *d*_*N*_ (or, as in Guill et al., 2021, the autotroph dispersal rate *d*_*A*_) is increased too far.

### 4.3 Oscillatory patterns create a positive eco-evo-spatial feedback loop

Our results suggest that self-organised pattern formation can generate a positive feedback loop between local biomass dynamics, adaptation of the trait distribution and regional dispersal that results in exceptionally high local diversity levels. Large trait variances, i.e., wide trait distributions, that result from the source-sink dynamics and (at times) disruptive local selection as described above, enable fast adaptation to the fluctuating local selection pressure, which leads to the formation of likewise asynchronous, i.e., spatially heterogeneous, oscillations of the community mean traits (Fig. 4c, d). In this state, mean traits are now differing between every pair of connected local communities for most of the time, not only between those exhibiting the defended and undefended local state. The dispersal of biomass of these different trait distributions between patches in turn enhances the positive effect of source-sink dynamics on local diversity. This translates into an increase of local trait variances, further enhances the flexibility of the trait distributions and thus the adaptive potential of the autotroph communities, closing the feedback loop with a spatial component. As the effectiveness of this eco-evo-spatial feedback loop depends on the strength of pattern formation, it promotes local diversity in highly connected metacommunities the most.

The enhancement of local diversity by the positive feedback loop is limited by two effects. First, fast adjustments of the trait distribution caused by high trait variances reduces the distance between the current mean trait value of a community and the momentary trait optimum, which leads to more stabilising, variance-reducing local selection again. Second, high flexibility of the trait distributions combined with strong source-sink dynamics drives convergence of the mean traits of undefended and defended communities. This results in a slight decrease of between-patch diversity and weakens the positive effect of source-sink dynamics on local diversity.

Feedbacks between ecological dynamics and the evolution of traits are increasingly recognised to be of high importance for the functioning of natural populations and communities in natural ecosystems. While they have already been extensively studied in simple predator-prey systems, we currently still lack insights for more complex (and thus more realistic) evolutionary food web- and meta-food web-models (M. Urban et al., 2008; Govaert et al., 2019). Our trophic metacommunity model contributes to fill this gap by highlighting an eco-evo-spatial feedback loop that gives rise to complex local and regional dynamics and relies on simultaneously taking into account the local trophic interactions, explicit nutrient dynamics, trait adaptation, and dispersal in metacommunities with realistic spatial structure. This emphasises that compiling several relevant ecological and evolutionary mechanisms into complex model systems can yield results which cannot be obtained from modelling these aspects separately. For example, in our metacommunity model, the continuous adaptation of the communities’ trait distributions are both cause and effect of high local functional diversity. Especially in the light of global land use change that affects species dispersal, it is of particular importance to assess the dispersal-driven eco-evolutionary feedbacks and their effect on diversity (Norberg et al., 2012; Govaert et al., 2019).

### 4.4 Relevance for ecological systems under global change

Self-organised pattern formation is a ubiquitous phenomenon emergent in many natural systems (Rietkerk and van de Koppel, 2008, Meron, 2015). Our results show that these patterns can enhance local diversity in realistically structured metacommunities across a wide range of size and connectance levels and thus highlight the relevance of this phenomenon for many different systems. As demonstrated, this diversity maintains the adaptability of autotroph communities to changes in their (a-)biotic environment (i.e., resource availability and predation pressure), which in our study where internally generated by the metacommunity dynamics. However, these changes may also be driven by external factors related to global change such as warming or eutrophication, which affect the fitness of autotrophs in a similar way (Binzer et al., 2012). High connectance of the surrounding metacommunity might therefore not only be of high importance for the local diversity of communities, but also for their adaptation to environmental changes that are caused or accelerated by climate change or intensified land use, e.g. further urbanisation or pesticide use (Balvanera et al., 2019; Chase et al., 2020). These same processes however may lead to loss and isolation of habitat patches in metacommunities, which results in lowered overall connectance (Chase et al., 2020). Through weakening the eco-evo-spatial feedback loop, loss of connectance could thus have disproportionately strong negative effects on local diversity of the remaining habitats (in line with Horváth et al., 2019) as well as reducing their adaptive potential to habitat change compared to healthy metacommunities.

In metacommunities with complex spatial structure, highly connected habitats (i.e., patches with high degree) are considered ecologically important as they influence and ensure rescue effects for a high number of other local habitats in the metacommunity (D. L. Urban and Keitt, 2001; Nakao and Mikhailov, 2010). As high-degree patches contribute above average to the overall connectance of a metacommunity, they are also vital for the positive effect of self-organised pattern formation on local functional diversity. From this perspective, they could therefore be discussed as ‘keystone’ patches or communities (Mouquet et al., 2013), though the impact of their removal was not specifically evaluated within the scope of this study.

### 4.5 Limitations & future perspectives

The homogenisation of local communities for very high dispersal rates always led to a complete loss of local and between-patch diversity even under strong pattern formation, showing that the described mechanisms for enhancing local diversity rely on the source-sink dynamics between the two local attractors. We did not observe maintenance of local and between patch diversity solely on the basis of fluctuating selection pressure, which may be related to the modelling assumption that the functional traits in each local community have a relatively rigid unimodal distribution. A reasonable step forward would therefore be to consider flexible trait distributions that often represents natural communities more accurately by allowing for multi-modality (Gaedke and Klauschies, 2017; Klauschies et al., 2018). The homogenisation process itself also merits further investigation, especially with the secondary diversity peak emerging in larger, sparsely connected metacommunities (Supporting Information, Appendix A.3). For example, it remains unclear whether there is a causal connection between the onset of pattern formation of the defended attractor and homogenisation of metacommunities.

In natural metacommunities, changes in size and connectance usually do not happen independent from each other, as assumed in this study. This limitation could be overcome by considering more specific case studies, where deletion of habitat patches also leads to the loss of its adjacent dispersal links or where a reduction of connectance leads to the loss of overall network connectivity and thus to the emergence of two or more smaller, but possibly on average better connected metacommunity components. In such a case, a loss of local diversity due to the smaller size could under pattern formation be compensated by higher internal connectance of the remaining, disconnected components.

### 4.6 Summary & Conclusion

In this study, we showed that self-organised pattern formation can considerably enhance the local functional diversity of autotroph communities embedded in metacommunities with realistic spatial structure through strengthening source-sink dynamics and reducing stabilising selection pressure. These mechanisms interact in a positive eco-evo-spatial feedback loop that is fueled by the emerging oscillatory patterns and is particularly strong in highly connected metacommunities, where local diversity could be maintained at high levels. As pattern formation is observed in many different natural systems, our findings are highly relevant in the light of anthropogenic habitat change that often destroys dispersal pathways, thereby increasing habitat isolation, lowering overall connectance of metacommunities and ultimately threatening the biodiversity in local habitats. Through potentially weakening complex feedback mechanisms, habitat isolation could endanger the diversity of local communities even more than previously assumed. We therefore advocate that emergent phenomena such as self-organised pattern formation should be included in the discussion about the sources and conservation of diversity in metacommunities subject to drivers of global change, and argue that it requires to jointly investigate the relevant ecological, evolutionary, and spatial mechanisms in complex model systems to obtain a comprehensive understanding of the processes.

## Acknowledgements

This study was supported by the German Research Foundation (DFG) under grant number GU 1645/2-1.

## A Supporting information

### A.1 Standard parameters

Model parametrisation follows the standard parameter values used in Guill et al. (2021). All simulations were run with the maximum attack rate (*a*_*max*_) of the heterotroph below a Hopf bifurcation of the undefended attractor so that the equilibrium of an isolated community was always a stable fixed point.

**Table A1:** Description and standard values of the model parameters. Adapted from Guill et al. (2021) with permission.

| parameter | description | dimension | value |
| --- | --- | --- | --- |
| $S$ | substrate supply concentration | $\text{mass} \cdot \text{area}^{-1}$ | 4.8 |
| $D$ | dilution rate | $\text{time}^{-1}$ | 0.3 |
| $N_H$ | half-saturation constant for nutrient uptake | $\text{mass} \cdot \text{area}^{-1}$ | 1.5 |
| $r_{max}$ | maximum growth rate of the autotroph | $\text{time}^{-1}$ | 1 |
| $r_{min}$ | minimum growth rate of the autotroph | $\text{time}^{-1}$ | 0.1 |
| $s$ | shape parameter of the growth rate function | - | 0.3 |
| $a_{max}$ | maximum attack rate of the heterotroph | $\frac{\text{area}}{\text{mass} \cdot \text{time}}$ | 1.3 |
| $b$ | scaling parameter of the attack rate function | - | 8 |
| $c$ | scaling parameter of the attack rate function | - | 0.4 |
| $h$ | handling time for the autotrophs | time | 0.53 |
| $e$ | conversion efficiency of the heterotroph | - | 0.33 |
| $\epsilon$ | scaling parameter of the boundary function | - | $10^{-3}$ |
| $\nu_{max}$ | maximum value of the trait variance | - | 0.25 |
| $d_N$ | diffusion coefficient of nutrients | - | varied |
| $d_A$ | diffusion coefficient of autotrophs | - | $10^{-5}$ |
| $d_H$ | diffusion coefficient of heterotrophs | - | $(10^{-4})$ |

### A.2 Observations and theoretical background of the relation between metacommunity structure and the strength of pattern formation

We calculated the long-term temporal variance of the autotroph biomass dynamics as a measure for the amplitude (or strength) of oscillatory self-organised patterns for all metacommunities that were used in the assessment of functional diversity (see Sections 2.3 and 2.4). Note that the patterns simultaneously manifest in the dynamics of the nutrients and heterotroph. Pattern amplitude increases with increasing metacommunity connectance (Figure A1a), best visible for high values of the nutrient diffusion coefficient *d*_*N*_. However, we observe a different effect over the size gradient: in metacommunities with low connectance, the pattern amplitude increases slightly from low to intermediate sizes, but only beyond the Turing instability threshold of the defended local attractor (Figure A1b), i.e. at very high values of *d*_*N*_ where the metacommunities already homogenise (see Section A.3 below). From intermediate to large sizes, pattern strength slightly declines. A similar observation can be made in fully connected metacommunities, but already below the instability threshold of the defended attractor. Beyond this secondary threshold, pattern strength however declines over the whole size range. The differences in how pattern strength changes over the size and connectance gradients might be caused by how the distribution of patch degrees, i.e., the number of dispersal links per habitat patch, in the networks changes over these gradients.

**Figure A1:**
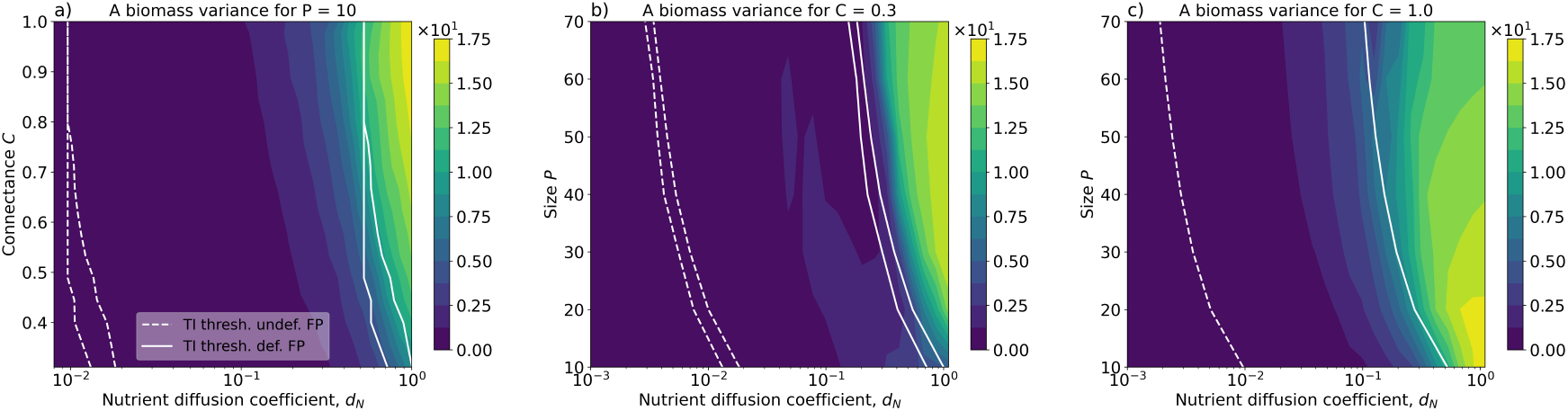
Long-term temporal variance of the autotroph biomass dynamics, averaged over patches and replicate metacommunities. Dynamics were evaluated after 100000 transient time steps.

In general, the amplitude of a developed pattern is determined by nonlinear interaction terms and their description thus requires a nonlinear stability analysis, e.g. in the form of amplitude equations (Banerjee et al., 2018; Chen and Buceta, 2019). This is often only possible close to the Turing instability thresholds. In our study, we are however interested in patterns that develop further away from the Turing instability threshold (in terms of the diffusion coefficient as bifurcation parameters). In scale-free networks, mean-field approximation can be used to estimate the pattern amplitude on a node, i.e., habitat patch (Nakao and Mikhailov, 2010; Mimar et al., 2019). For a defined mean field, the pattern amplitude on a patch is proportional to the diffusion coefficients and the respective patch degree. While the random geometric graphs that form the basis of our metacommunity networks are not scale-free, we also observe an increase of the pattern amplitude when changing the diffusion coefficient *d*_*X*_ in such a way that dynamics are led further into the Turing instability region (Box 1, Figure A1). Similar observations were made in previous studies (Hata et al., 2014; Guill et al., 2021). With increasing size and connectance, the patch degrees increase (Figure 1). However, modifying the underlying network structure also alters the respective mean field and therefore it is not possible to deduce a direct relation between the patch degree of nodes and the resulting pattern amplitude in different metacommunity networks.

Importantly, increasing size or connectance not only increases the degrees of individual patches, but also alters the degree distribution of the entire network in a distinct way. While in random geometric graphs low connectance is associated with a degree distribution constrained to low patch degrees (right-skewed), at intermediate connectance the degree distribution is wide and symmetrical (Figure A2). At high connectance the degree distribution is again more constrained, now to high patch degrees (left-skewed), until in fully connected networks all patches have the same degree *P* − 1. This skew and narrowing of the degree distribution occurs while the *d*_*N*_ threshold for the Turing instability remains constant once the first patch in the metacommunity reaches the maximum degree of *P* − 1 (which is very likely to occur in networks with *C* ≥ 0.71). Conversely, by increasing the size of a metacommunity (with low connectance), the degree distribution is always largely symmetrical and gradually widens (Figure A3). Additionally, the *d*_*N*_ threshold value for the Turing instability keeps changing with size as subsequently larger maximum degrees are reached. Analysing scale-free and Erd?os-Rényi random graphs, Mimar et al. (2019) found that the degree distribution affects the amplitude of the emerging patterns mostly when it is wide (i.e., when connectance is not too high), but not in highly connected networks with narrow degree distribution. This might indicate that a wide degree distribution limits the amplitude of a developing pattern even if the patch degrees themselves are increasing. Considering that the topology of patch networks embedded in physical space very likely differs from that of the stochastic network models used by Mimar et al. (2019), additional work is however needed to provide a systematic insight into the causal relation between different structural metrics of metacommunities and the amplitude of developed patterns.

**Figure A2:**
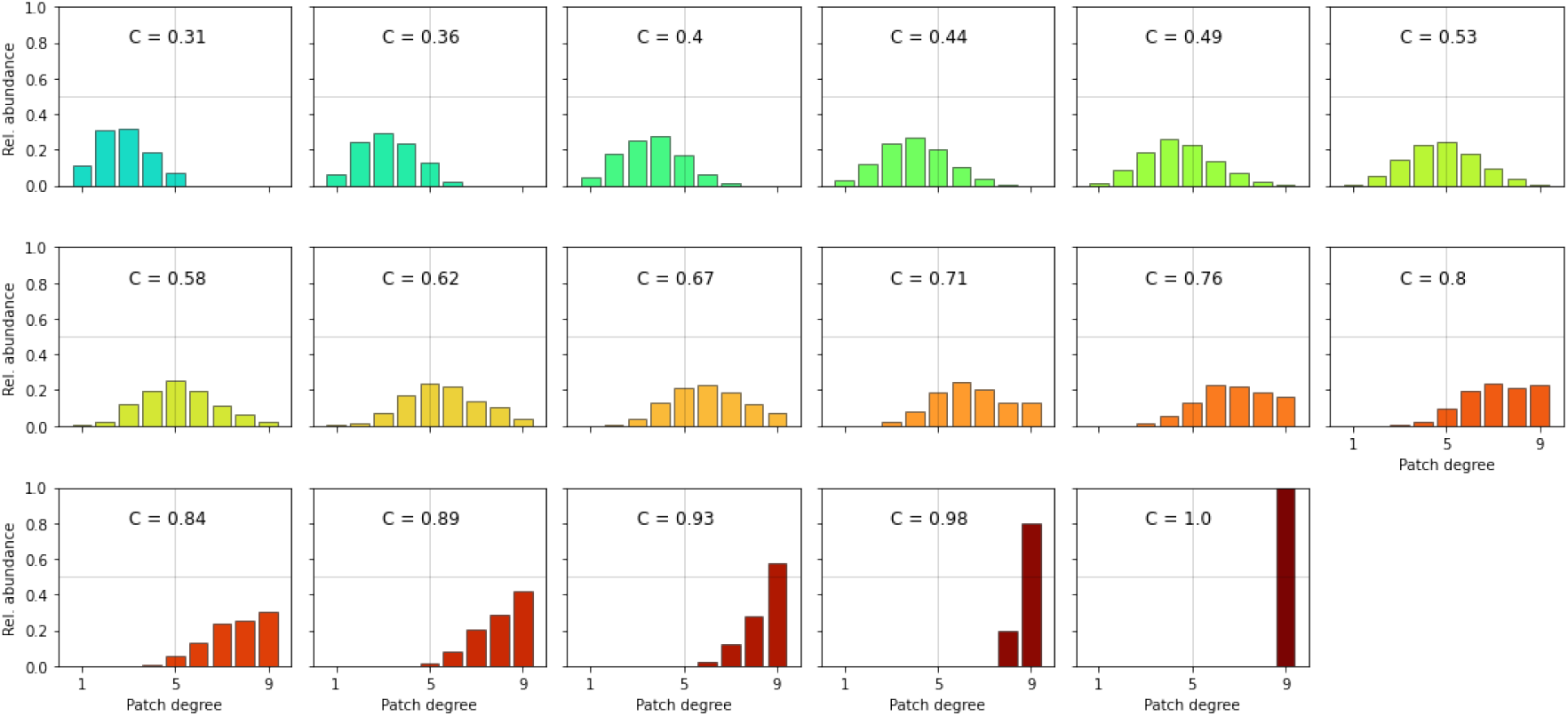
Change of degree distribution with connectance in 100 networks of P = 10 patches

**Figure A3:**
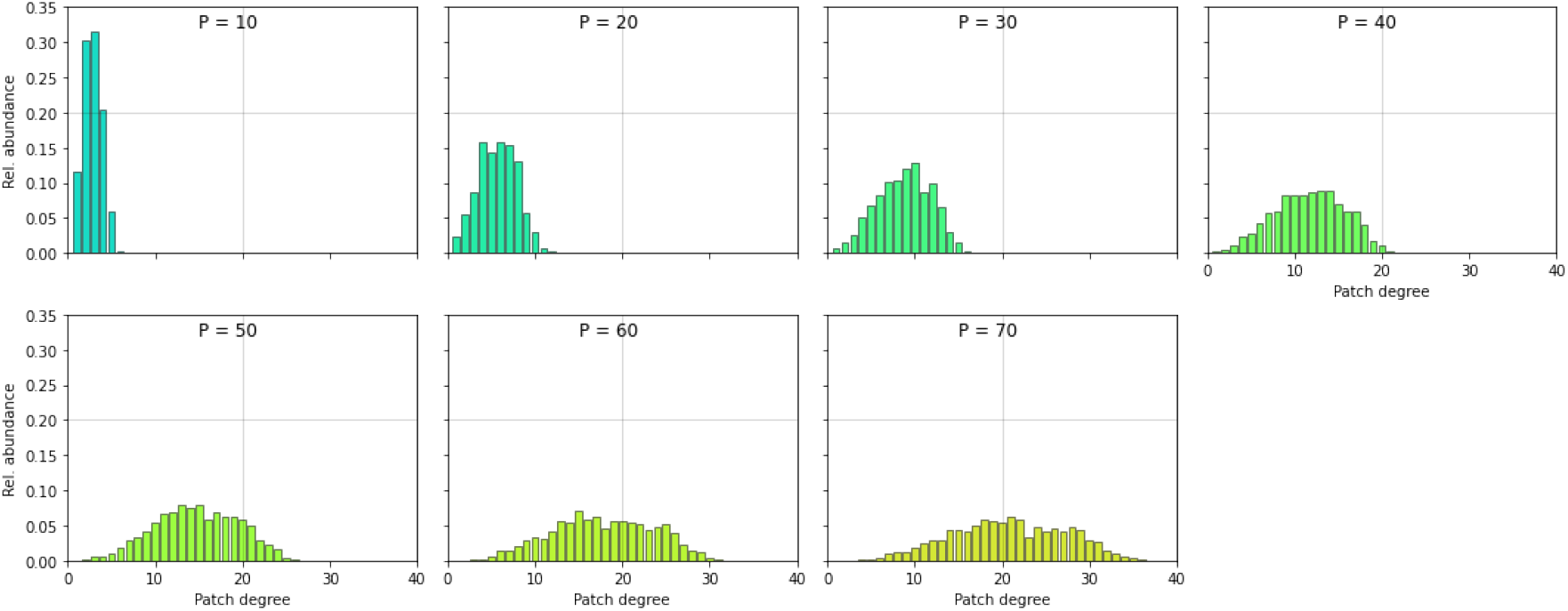
Change of degree distribution with size in 40 networks of connectance C = 0.3

### A.3 Homogenisation at high nutrient diffusion

We observed homogenisation in the metacommunities, where all patches exhibit a common (defended or undefended) state rather than initialisation-dependent local bistability (see section 2.1). The homogeneous metacommunities still exhibited oscillatory pattern formation with large amplitudes (Figure A1), but due to the sudden loss of *β*-diversity, 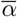-diversity swiftly decreased towards zero in every homogeneous metacommunity (Figure 3). Homogenisation occurred for metacommunities of all tested sizes and connectance levels for high values of the nutrient diffusion coefficient *d*_*N*_ in proximity to the Turing instability threshold of the defended local attractor (the instability threshold of the undefended local attractor is crossed at lower *d*_*N*_). However, not all metacommunities homogenised at the same *d*_*N*_ and towards the same attractor (Figure A4), which points towards a multistability between the locally bistable and homogenised states that might be dependent on initialisation and the underlying metacommunity structure. With increasing *d*_*N*_, first a rising fraction of metacommunities homogenises on a common defended state, but then the ho-mogenisation regime shifts to a common undefended state until defended communities both in bistable and homogeneous metacommunities disappear altogether.

Only in larger(*>* 40*P*), weakly connected metacommunities, a window of local bistability between homogenisation on the common defended and undefended attractor occurs (Figure A4a). There, some of the metacommunities that homogenised on a common defended state at lower *d*_*N*_ now (at higher *d*_*N*_) maintain both undefended and defended communities again. This might coincide with the Turing instability of the defended communities. Within this window, 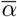- and *β*-diversity are maintained at nonzero levels again, resulting in the secondary diversity peaks visible in Figure 3c and d. Bistable metacommunities in this state show 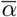-diversity levels that are higher than the peak levels observed for metacommunities of same size and connectance but for *d*_*N*_ values below the initial homogenisation, which might be caused by the exceptionally large pattern amplitudes (see Discussion for details). An even further increase of *d*_*N*_ however ultimately leads to homogenisation on a common undefended attractor and again loss of *β*-as well as 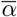-diversity.

In fully connected metacommunities of any size, homogenisation sets in more swiftly and a secondary window of bistability does not occur (Figures 3e and A4b). It could thus be possible that the wide degree distribution in large, sparsely connected metacommunities and especially the presence of patches with few dispersal links (low degree) or relatively isolated patch groups can delay the homogenisation within metacommunities. Complex interactions between metacommunity structure and dynamics might become only visible when the studied structures are large enough (Nakao and Mikhailov, 2010).

**Figure A4:**
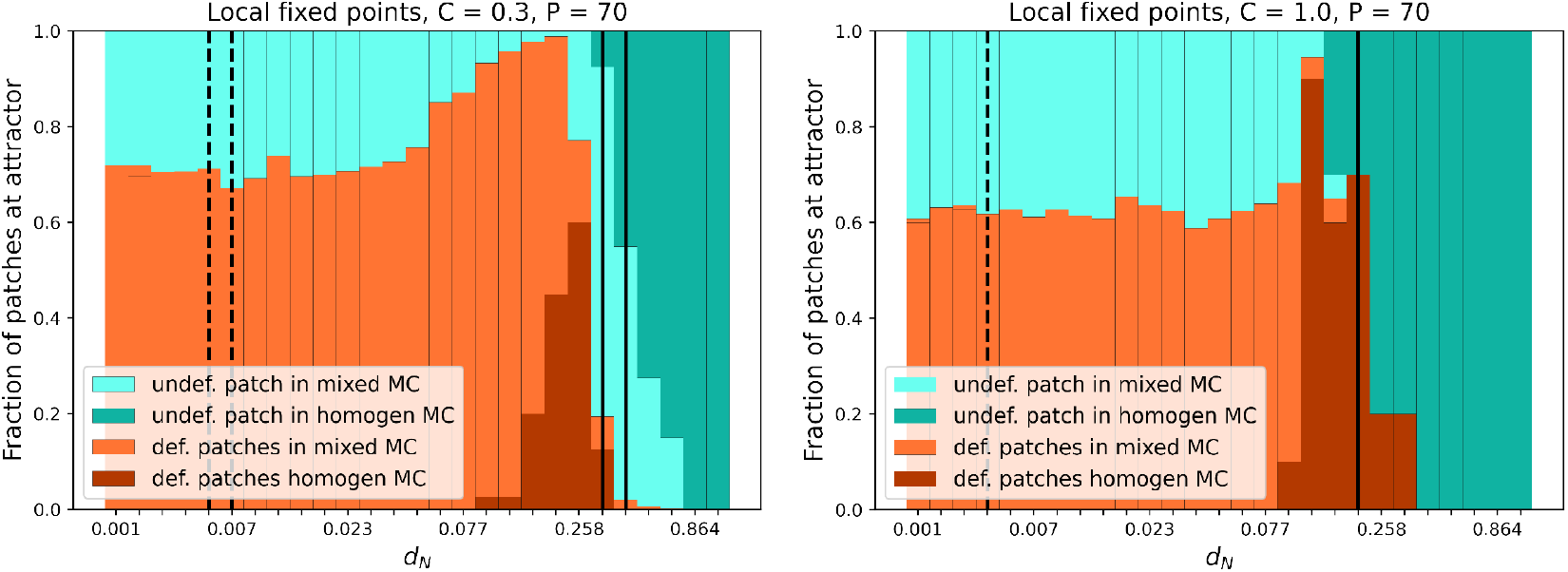
Attractor distribution and metacommunity homogenisation across nutrient mobility gradient. Occurrence frequency of defended and undefended patches in metacommunities where both attractors were present(’mixed’) as well as occurrence frequency of defended and undefended patches in homogenised metacommunities, where all patches converged towards the respective local attractor. The category “undefended” was defined as mean trait ≥0.7,“defended“ as mean trait < 0.7. Assessed from long-term temporal averages of 40 metacommunities.

## Notes

### Competing Interest Statement

The authors have declared no competing interest.

## References

Amarasekare, Priyanga (2003). “Competitive coexistence in spatially structured environments: a synthesis”. Ecol. Lett. 6, pp. 1109–1122.

Aufderheide, Helge E (2014). “Implications of eigenvector localization for dynamics on complex networks”. Dissertation. Technische Universität Dresden.

Balvanera, P., A. Pfaff, A. Viña, E. García-Frapolli, L. Merino, P. A. Minang, N. Nagabhatla, S. A. Hussain, and A. A. Sidorovich (2019). “Chapter 2.1 Status and Trends - Drivers of Change. In: Global assessment report of the Intergovernmental Science-Policy Platform on Biodiversity and Ecosystem Services”. In: ed. by E. S. Brondízio, J. Settele, S. Díaz, and H. T. Ngo. Bonn, Germany, p. 152.

Banerjee, Malay, S Ghorai, and Nayana Mukherjee (2018). “Study of cross-diffusion induced Turing patterns in a ratio-dependent prey-predator model via amplitude equations”. en. Applied Mathematical Modelling 55, pp. 383–399.

Bauer, Barbara, Michael Kleyer, Dirk C. Albach, Bernd Blasius, Ulrich Brose, Thalita Ferreira-Arruda, Ulrike Feudel, Gabriele Gerlach, Christian Hof, Holger Kreft, Lucie Kuczynski, Kertu Lõhmus, Stefanie Moorthi, Christoph Scherber, Stefan Scheu, Gerhard Zotz, and Helmut Hillebrand (2021). “Functional trait dimensions of trophic metacommunities”. Ecography 44.10, pp. 1486–1500.

Bennett, Jamie J R, Anabele S Gomes, Michel A Ferré, Bidesh K Bera, Fabian Borghetti, Ragan M Callaway, and Ehud Meron (2023). “Evidence for scale-dependent root-antation feedback and its role in halting the spread of a pantropical shrub into an endemic sedge”. en. PNAS Nexus 2.1. Ed. by Alberto Guadagnini, pgac294.

Binzer, Amrei, Christian Guill, Ulrich Brose, and Björn C. Rall (2012). “The dynamics of food chains under climate change and nutrient enrichment”. en. Philosophical Transactions of the Royal Society B: Biological Sciences 367.1605, pp. 2935–2944.

Bornholdt, Stefan and Heinz Georg Schuster (2001). Handbook of graphs and networks: From the Genome to the Internet. Wiley Online Library.

Brechtel, Andreas, Philipp Gramlich, Daniel Ritterskamp, Barbara Drossel, and Thilo Gross (2018). “Master stability functions reveal diffusion-driven pattern formation in networks”. Phys. Rev. E 97.032307.

Brown, James H. and Astrid Kodric-Brown (1977). “Turnover rates in insular biogeography: effect of immigration on extinction”. Ecology 58.2, pp. 445–449.

Cadotte, Marc W. (2006). “Dispersal and species diversity: a meta-analysis”. The American Naturalist 167.6, pp. 913–924.

Chase, Jonathan M., Alienor Jeliazkov, Emma Ladouceur, and Duarte S. Viana (2020). “Biodiversity conservation through the lens of metacommunity ecology”. en. Annals of the New York Academy of Sciences 1469.1, pp. 86–104.

Chen, Yanyan and Javier Buceta (2019). “A non-linear analysis of Turing pattern formation”. en. PLOS ONE 14.8. Ed. by Duccio Fanelli, e0220994.

Cornacchia, Loreta, Daphne van der Waal, Geraldene Wharton, Sara Puijalon, and Tjeerd J. Bouma (2018). “Landscapes of facilitation: how self-organized patchiness of aquatic macrophytes promotes diversity in streams”. Ecology 99, pp. 832–847.

Eigentler, L and J A Sherratt (2020). “Spatial self-organisation enables species coexistence in a model for savanna ecosystems”. J. Theor. Biol. 487, p. 110122.

Eigentler, L. (2021). “Species coexistence in resource-limited patterned ecosystems is facilitated by the interplay of spatial self-organisation and intraspecific competition”. Oikos 130, pp. 609–623.

Foley, Jonathan A., Ruth DeFries, Gregory P. Asner, Carol Barford, Gordon Bonan, Stephen R. Carpenter, F. Stuart Chapin, Michael T. Coe, Gretchen C. Daily, Holly K. Gibbs, Joseph H. Helkowski, Tracey Holloway, Erica A. Howard, Christopher J. Kucharik, Chad Monfreda, Jonathan A. Patz, I. Colin Prentice, Navin Ramankutty, and Peter K. Snyder (2005). “Global Consequences of Land Use”. en. Science 309.5734, pp. 570–574.

Gaedke, Ursula and Toni Klauschies (2017). “Analyzing the shape of observed trait distributions enables a data-based moment closure of aggregate models”. Limnol. Oceanog. Meth. 15.12, pp. 979–994.

Galassi, Mark, Jim Davies, James Theiler, Brian Gough, Gerard Jungman, Patrick Alken, Michael Booth, Fabrice Rossi, and Rhys Ulerich (2002). GNU scientific library. Network Theory Limited.

Govaert, Lynn, Emanuel A. Fronhofer, Sébastien Lion, Christophe Eizaguirre, Dries Bonte, Martijn Egas, Andrew P. Hendry, Ayana De Brito Martins Carlos J. Melián, Joost A. M. Raeymaekers, Irja I. Ratikainen, Bernt-Erik Saether, Jennifer A. Schweitzer, and Blake Matthews (2019). “Eco-evolutionary feedbacks—Theoretical models and perspectives”. en. Functional Ecology 33.1. Ed. by Charles Fox, pp. 13–30.

Guill, Christian, Janne Hülsemann, and Toni Klauschies (2021). “Self-organised pattern formation increases local diversity in metacommunities”. Ecology letters 24.12, pp. 2624–2634.

Haegeman, Bart and Michel Loreau (2014). “General relationships between consumer dispersal, resource dispersal and metacommunity diversity”. Ecology Letters 17.2, pp. 175–184.

Harris, Charles R., K. Jarrod Millman, Stéfan J. Van Der Walt, Ralf Gommers, Pauli Virtanen, David Cournapeau, Eric Wieser, Julian Taylor, Sebastian Berg, Nathaniel J. Smith, Robert Kern, Matti Picus, Stephan Hoyer, Marten H. van Kerkwijk, Matthew Brett, Allan Haldane, Jaime Fernández del Río, Mark Wiebe, Pearu Peterson, Pierre Gérard-Marchant, Kevin Sheppard, Tyler Reddy, Warren Weckesser, Hameer Abbasi, Christoph Gohlke, and Travis E. Oliphant (2020). “Array programming with NumPy”. Nature 585.7825, pp. 357–362.

Hata, Shigefumi, Hiroya Nakao, and Alexander S. Mikhailov (2014). “Dispersal-induced destabilization of metapopulations and oscillatory Turing patterns in ecological networks”. en. Scientific Reports 4.1, p. 3585.

Hedetniemi, Stephen T., David P. Jacobs, and Vilmar Trevisan (2016). “Domination number and Laplacian eigenvalue distribution”. European Journal of Combinatorics 53, pp. 66–71.

Hindmarsh, Alan C., Peter N. Brown, Keith E. Grant, Steven L. Lee, Radu Serban, Dan E. Shumaker, and Carol S. Woodward (2005). “SUNDIALS: Suite of nonlinear and differential/algebraic equation solvers”. ACM Trans. Math. Software (TOMS) 31.3, pp. 363–396.

Holland, Matthew D. and Alan Hastings (2008). “Strong effect of dispersal network structure on ecological dynamics”. Nature 456.7223, p. 792.

Holyoak, Marcel, Mathew A Leibold, and Robert D Holt (2005). Metacommunities: spatial dynamics and ecological communities. University of Chicago Press.

Horváth, Zsófia, Robert Ptacnik, Csaba F Vad, and Jonathan M Chase (2019). “Habitat loss over six decades accelerates regional and local biodiversity loss via changing landscape connectance”. Ecology letters 22.6, pp. 1019–1027.

Hunter, John D. (2007). “Matplotlib: A 2D graphics environment”. Comput. Science & Eng. 9.3, p. 90.

Joblib Development Team (2021). Joblib: running Python functions as pipeline jobs.

Kéfi, Sonia, Maarten B. Eppinga, Peter C. de Ruiter, and Max Rietkerk (2010). “Bistability and regular spatial patterns in arid ecosystems”. Theor. Ecol. 3.4, pp. 257–269.

Klauschies, Toni, Renato M. Coutinho, and Ursula Gaedke (2018). “A beta distribution-based moment closure enhances the reliability of trait-based aggregate models for natural populations and communities”. Ecol. Mod. 381, pp. 46–77.

LeCraw, Robin M., Diane S. Srivastava, and Gustavo Q. Romero (2014). “Metacommunity size influences aquatic community composition in a natural mesocosm landscape”. en. Oikos 123.8, pp. 903–911.

Leibold, Mathew A., Marcel Holyoak, Nicolas Mouquet, Priyanga Amarasekare, Jonathan M. Chase, Martha F. Hoopes, Robert D. Holt, Jonathan B. Shurin, Richard Law, David Tilman,M. Loreau, and A. Gonzalez (2004). “The metacommunity concept: a framework for multi-scale community ecology”. Ecol. Lett. 7.7, pp. 601–613.

Liu, Quan-Xing, Peter M.J. Herman, Wolf M. Mooij, Jef Huisman, Marten Scheffer, Han Olff, and Johan Van De Koppel (2014). “Pattern formation at multiple spatial scales drives the resilience of mussel bed ecosystems”. Nature communications 5.1, pp. 1–7.

Logue, Jürg B., Nicolas Mouquet, Hannes Peter, and Helmut Hillebrand (2011). “Empirical approaches to metacommunities: a review and comparison with theory”. en. Trends in Ecology & Evolution 26.9, pp. 482–491.

Malchow, Horst (1993). “Spatio-temporal pattern formation in nonlinear non-equilibrium plankton dynamics”. Proc. R. Soc. Lond. B 251.1331, pp. 103–109.

Meron, Ehud (2015). Nonlinear physics of ecosystems. CRC Press Boca Raton, FL.

Mimar, Sayat, Mariamo Mussa Juane, Juyong Park, Alberto P. Muñuzuri, and Gourab Ghoshal (2019). “Turing patterns mediated by network topology in homogeneous active systems”. en. Physical Review E 99.6, p. 062303.

Mouquet, Nicolas, Dominique Gravel, François Massol, and Vincent Calcagno (2013). “Extending the concept of keystone species to communities and ecosystems”. Ecology Letters 16.1, pp. 1–8.

Mouquet, Nicolas and Michel Loreau (2003). “Community patterns in source-sink metacommunities”. Am. Nat. 162.5, pp. 544–557.

Murray, James D., ed. (2003). Mathematical Biology: II: Spatial Models and Biomedical Applications. Third Edition. Vol. 18. Springer eBook Collection Mathematics and Statistics. New York, NY: Springer New York.

Nakao, Hiroya and Alexander S. Mikhailov (2010). “Turing patterns in network-organized activator– inhibitor systems”. Nature Physics 6.7, pp. 544–550.

Nathan, J., E. Meron, and J. von Hardenberg (2013). “Spatial instabilities untie the exclusion-principle constraint on species coexistence”. J. Theor. Biol. 335, pp. 198–204.

Newbold, Tim, Lawrence N. Hudson, Samantha L. L. Hill, Sara Contu, Igor Lysenko, Rebecca A. Senior, Luca Börger, Dominic J. Bennett, Argyrios Choimes, Ben Collen, Julie Day, Adriana De Palma, Sandra Díaz, Susy Echeverria-Londonõ, Melanie J. Edgar, Anat Feldman, Morgan Garon, Michelle L. K. Harrison, Tamera Alhusseini, Daniel J. Ingram, Yuval Itescu, Jens Kattge, Victoria Kemp, Lucinda Kirkpatrick, Michael Kleyer, David Laginha Pinto Correia, Callum D. Martin, Shai Meiri, Maria Novosolov, Yuan Pan, Helen R. P. Phillips, Drew W. Purves, Alexandra Robinson, Jake Simpson, Sean L. Tuck, Evan Weiher, Hannah J. White, Robert M. Ewers, Georgina M. Mace, Jörn P. W. Scharlemann, and Andy Purvis (2015). “Global effects of land use on local terrestrial biodiversity”. en. Nature 520.7545, pp. 45–50.

Norberg, Jon, Dennis P. Swaney, Jonathan Dushoff, Juan Lin, Renato Casagrandi, and Simon A. Levin (2001). “Phenotypic diversity and ecosystem functioning in changing environments: a theoretical framework”. Proc. Nat. Acad. Sci. 98.20, pp. 11376–11381.

Norberg, Jon, Mark C. Urban, Mark Vellend, Christopher A. Klausmeier, and Nicolas Loeuille (2012). “Eco-evolutionary responses of biodiversity to climate change”. en. Nature Climate Change 2.10, pp. 747–751.

Othmer, Hans G. and L.E. Scriven (1971). “Instability and dynamic pattern in cellular networks”. J. Theor. Biol. 32.3, pp. 507–537.

Rietkerk, Max and Johan van de Koppel (2008). “Regular pattern formation in real ecosystems”. Trends Ecol. Evol. 23.3, pp. 169–175.

Riva, Federico, Caio Graco-Roza, Gergana N. Daskalova, Emma J. Hudgins, Jayme M. M. Lewthwaite, Erica A. Newman, Masahiro Ryo, and Stefano Mammola (2023). “Toward a cohesive understanding of ecological complexity”. en. Science Advances 9.25, eabq4207.

Ryser, Remo, Johanna Häussler, Markus Stark, Ulrich Brose, Björn C. Rall, and (2019). “The biggest losers: habitat isolation deconstructs complex food webs from top to bottom”. Proc. R. Soc. Lond. B 286.1908, p. 20191177.

Staddon, Philip, Zöe Lindo, Peter D. Crittenden, Francis Gilbert, and Andrew Gonzalez (2010). “Connectivity, non-random extinction and ecosystem function in experimental metacommunities: Ecosystem function in experimental metacommunities”. en. Ecology Letters 13.5, pp. 543–552.

Tews, J., U. Brose, V. Grimm, K. Tielbörger, M. C. Wichmann, M. Schwager, and F. Jeltsch (2004). “Animal species diversity driven by habitat heterogeneity/diversity: the importance of keystone structures”. Journal of Biogeography 31.1, pp. 79–92.

Thompson, Patrick L, Bronwyn Rayfield, and Andrew Gonzalez (2017). “Loss of habitat and connectivity erodes species diversity, ecosystem functioning, and stability in metacommunity networks”. Ecography 40.1, pp. 98–108.

Tirok, Katrin, Barbara Bauer, Kai Wirtz, and Ursula Gaedke (2011). “Predator-prey dynamics driven by feedback between functionally diverse trophic levels”. PloS One 6.11, e27357.

Turing, Alan Mathison (1952). “The chemical basis of morphogenesis”. Phil. Trans. R. Soc. Lond. B 237.641, pp. 37–72.

Urban, Dean L. and Timothy Keitt (2001). “Landscape connectivity: a graph-theoretic perspective”. Ecology 82.5, pp. 1205–1218.

Urban, M., M. Leibold, P. Amarasekare, L. Demeester, R. Gomulkiewicz, M. Hochberg, C. Klausmeier, N. Loeuille, C. Demazancourt, and J. Norberg (2008). “The evolutionary ecology of metacommunities”. en. Trends in Ecology & Evolution 23.6, pp. 311–317.

Van Der Kolk, Jasper, Guillermo García-Pérez, Nikos E. Kouvaris, M. Ángeles Serrano, and Marián Boguñá (2023). “Emergence of Geometric Turing Patterns in Complex Networks”. en. Physical Review X 13.2, p. 021038.

van der Valk, Arnold G. and Barry G. Warner (2009). “The development of patterned mosaic landscapes: an overview”. Plant Ecology 200.1, pp. 1–7.

Virtanen, Pauli, Ralf Gommers, Travis E. Oliphant, Matt Haberland, Tyler Reddy, David Cournapeau, Evgeni Burovski, Pearu Peterson, Warren Weckesser, Jonathan Bright, Stéfan J. van der Walt, Matthew Brett, Joshua Wilson, K. Jarrod Millman, Nikolay Mayorov, Andrew R. J. Nelson, Eric Jones, Robert Kern, Eric Larson, C. J. Carey, İlhan Polat, Yu Feng, Eric W. Moore, Jake Van der Plas, Denis Laxalde, Josef Perktold, Robert Cimrman, Ian Henriksen, E. A. Quintero, Charles R. Harris, Anne M. Ar chibald, Antônio H. Ribeiro, Fabian Pedregosa, Paul van Mulbregt, and SciPy 1.0 Contributors (2020). “SciPy 1.0: fundamental algorithms for scientific computing in Python”. Nature methods 17.3, pp. 261–272.

Wirtz, Kai and Bruno Eckhardt (1996). “Effective variables in ecosystem models with an applica-tion to phytoplankton succession”. Ecol. Mod. 92.1, pp. 33–53.

